# Acquisition, co-option, and duplication of the *rtx* toxin system and the emergence of virulence in *Kingella*

**DOI:** 10.1101/2022.11.28.518221

**Authors:** Daniel P. Morreale, Eric A. Porsch, Brad K. Kern, Joseph W. St Geme, Paul J. Planet

## Abstract

The *Kingella* genus includes two pathogenic species, namely *K. kingae* and *K. negevensis*, as well as strictly commensal species. Both *K. kingae* and *K. negevensis* secrete a toxin called RtxA that is absent in the commensal species. Phylogenetic analysis demonstrates that the toxin-encoding operon *rtxCrtxAtolC* was acquired by a common ancestor of the pathogenic *Kingella* species and that a preexisting type I secretion system was co-opted for toxin export. Subsequent genomic reorganization distributed the toxin machinery across two loci, with 30-35% of *K. kingae* strains containing two copies of the *rtxA* toxin gene. The *rtxA* duplication is largely clonal and strongly associated with invasive disease. In assays with isogenic strains, a single copy of *rtxA* was associated with reduced virulence *in vitro*. This study establishes the critical steps in the evolutionary transition from commensal to pathogen, including horizontal gene transfer, co-option of an existing secretion system, and gene duplication.

## Introduction

*Kingella kingae* is a Gram-negative coccobacillus that was originally thought to be a rare cause of human disease and is now known to be an important pathogen, especially in young children. *K. kingae* initiates infection by colonizing the oropharynx, where it can persist for weeks to months without causing symptoms.^1–5^ On occasion, *K. kingae* translocates across the oropharyngeal epithelial barrier, enters the bloodstream, and disseminates to distant sites such as bones, joints, and the endocardium to cause disease, generally in the setting of viral infection of the oral cavity or upper respiratory tract.^1^ In many parts of the world, *K. kingae* has emerged as the most common cause of septic arthritis in children between 6 and 48 months of age.^6^

The development of invasive disease by *K. kingae* is facilitated by several virulence factors, including an RTX family toxin, type IV pili, an autotransporter adhesin, a lipopolysaccharide-associated exopolysaccharide, and a polysaccharide capsule.^7^ Together, these factors are thought to allow *K. kingae* to adhere to and lyse host cells, resist complement-mediated killing and neutrophil phagocytosis in the bloodstream, and subvert neutrophil killing at the site of disease. Of the five species that comprise the *Kingella* genus, only *K. kingae* and *K. negevensis* have been repeatedly associated with invasive disease in immunocompetent individuals.^5,8^ These two species are also distinguished from the rest of the genus by the presence of an RTX family toxin called RtxA, which is encoded by the *rtxA* gene and causes β-hemolysis and toxicity to a variety of human cell types.^9^ In both *K. kingae* and *K. negevensis*, the toxin-associated genes are hypothesized to have been acquired as a part of the mobile genetic element, though no study has defined the evolutionary history of this locus in detail in diverse *K. kingae* isolates.^9–11^

The RtxA toxin is highly conserved among *K. kingae* isolates and is required for morbidity and mortality in an animal model of invasive disease.^12^ RtxA is modified by the RtxC acyltransferase and is secreted from the organism by a type I secretion system (TISS) that is encoded by the products of the *rtxB, rtxD*, and *tolC* genes. The toxin-associated genes are present at two loci in *K. kingae*, including one locus that contains *rtxB, rtxD*, and *rtxC* and a second locus that contains *rtxC, rtxA*, and the *tolC* gene, which encodes the TolC outer membrane protein.^10^ In at least one strain (KWG-1), there are two copies of both *rtxA* and *tolC*, with the second copy of these genes downstream of *rtxB* and *rtxD*.^7,10,13^ The impact of this duplication on virulence and pathogenicity is unknown.

Prior studies have shown that certain clonal groups identified by pulsed-field gel electrophoresis (PFGE) or multilocus sequence typing (MLST) are associated with invasive disease. For example, 70% of *K. kingae* invasive isolates belong to PFGE groups B, H, K, N, and P (out of 32 total groups). PFGE group K has been associated with occult bacteremia, group N with skeletal infections, and group P with endocarditis.^14^ Similarly, ST-6, ST-14, ST-23, ST-25, and ST-66 have been implicated in *K. kingae* outbreaks in daycare centers.^3,15,16^ No detailed analysis has been conducted on *K. kingae* genomes to identify virulence-associated genes or genotypes in these sequence types.

In this study, we hypothesized that there are specific genomic features that are responsible for both the emergence of virulence in *K. kingae* and *K. negevensis* and the association of particular lineages with a greater likelihood of invasiveness. We found that the efficient secretion of RtxA is the result of co-option of a preexisting type I secretion system, a critical step in the evolution of *K. kingae* and *K. negevensis* virulence. Phylogenetic analysis of whole genomes from 88 *K. kingae* clinical isolates sequenced for this study along with all 52 publicly available *K. kingae* genomes allowed us to test for lineages and genomic features associated with invasive disease. We discovered that a duplication of *rtxA* is associated with invasive disease and is concentrated primarily in a single clade. Additional analysis established that a single copy of *rtxA* is associated with reduced virulence compared to two copies of *rtxA in vitro*.

## Material and Methods

### *In silico* analysis of diverse *Kingella* isolates

#### Whole Genome Sequencing

Isolates used in this study and all available metadata are listed in Table S1. Clinical isolates for sequencing (88 isolates, primarily recovered from patients in Israel) and genotyping (an additional 131 global isolates) were selected from a large collection of more than 500 *K. kingae* isolates collected from the early 1990’s. Additionally, the sequencing reads from 52 publicly available genomes were downloaded from the NCBI Sequence Read Archive (SRA) database on August 5, 2020. Publicly available genomes include isolates from as early as 1966. Genomic DNA (gDNA) for sequencing and genotyping was extracted using the Wizard Genomic DNA purification kit (Promega) as directed for Gram-negative bacterial species. The gDNA concentration was determined via NanoDrop 2000 (Thermo Scientific).

Extracted gDNA was fragmented and barcoded using the Nextera XT kit (Illumina), following the manufacturer’s instructions. Genomes were sequenced on an Illumina HiSeq 2500 machine using the 100SR single end read protocol. Sequencing adapters and low-quality ends with a PHRED score less than 15 were trimmed from reads using Trim Galore v. 0.6.4.^17^ Trimmed reads and reads downloaded from the SRA databases were assembled using SPAdes assembler v. 3.14.0 and default settings for single end Illumina reads.^18,19^ Quality control data for both trimming and assembly are shown in Table S2 and was calculated using QUAST. Assembled scaffolds were annotated using Prokka v. 1.14.6, using the following flags and options: --addgenes --kingdom Bacteria --genus *Kingella* --species *kingae*.^20^ Finally, nucleotide and amino acid FASTA files generated as an output from Prokka for all isolates were compiled into two databases for local blast searches. All reads and assemblies are available through SRA (BioProject PRJNA896475). To validate that all strains were members of the species *K. kingae*, the average nucleotide identity score (ANI) was calculated for each isolate relative to the isolate *K. kingae* strain KWG-1 with FastANI v.1.32.^21^ Additionally, FastANI was used to generate an output matrix later plotted as a heatmap in R v.4.1.2. Quast v. 5.0.2 was used to determine the quality of all assemblies (data summarized in Table S1 and Table S2), using *K. kingae* KWG-1 as the reference sequence.^22^ Sequence type was assigned using the *K. kingae* PasteurMLST scheme.^15,22,23^

### Long read sequencing of K. negevensis strains

*K. negevensis* gDNA was extracted using the Wizard HMW DNA extraction Kit (Promega) following the manufacturer’s instructions. Following extraction, DNA quality was determined using TapeStation Genomic DNA Screentape (Agilent) and quantified using a NanoDrop 2000 (Thermo Scientific). For sequencing, libraries were generated using the Rapid Barcoding Sequencing Kit (SQK-RBK004, Oxford Nanopore Technologies), using 400 ng of template DNA, and following the manufacturer’s instructions. Sequencing was performed on the MinION Mk-1B using the Spot-ON Flow Cell R9 (Lot 11002153, Oxford Nanopore Technologies). Base calling was performed using the MinKNOW software with increased stringency. Low quality reads and reads less than 1 kB in length were removed with FiltLong. Read pile-ups were conducted using pooled reads from all isolates against the *K. kingae* KWG-1 genome using SAMtools and visualized with the R packages Minimap2 and Sushi.^24–26^ *De novo* assemblies of each isolate were performed with the Raven genome assembly tool, and copy number was determined using BLASTn.^27^

### Pangenome characterization

The GFF output file from Prokka was analyzed with Roary v.3.13.0 to determine genes in the core and pangenome.^28^ Roary was run with the following options selected: create multiFASTA alignments of core genes using PRANK (-e), generate a fast core gene alignment with MAFFT^29^ (-n), at least 95% percent of isolates must encode a gene for the gene to be considered a core gene (-cd 95), generate R plots (-r), and cluster paralogs (-s). All other settings were default. To identify known virulence factors in the core and pangenome, assembled contigs were analyzed against the VFDB and MEGARES database with Abricate v.1.0.1 and NCBI AMRFinder v.3.9.3, using default settings.^30–33^ Scoary v.1.6.16 was used to perform genome wide association studies to identify genes associated with invasive isolates, relative to carrier isolates, with default settings.^34^

### Phylogenetic reconstructions

Single nucleotide polymorphisms (SNPs) were identified in trimmed sequencing reads from all stains that passed quality control cutoffs with Snippy v. 4.6.0 by generating a whole genome alignment against *K. kingae* KWG-1 as a reference (Genbank file downloaded from NCBI September 01, 2020).^35^ The output alignment file was cleaned using Snippy clean module, and a phylogenetic tree was reconstructed in RAxML v.8.2.4, along with 100 bootstraps, the GTR substitution model, GAMMA model of rate heterogeneity, rapid hill climbing (-f d), and a random number seed of 1977.^36^ These settings were used for all nucleotide trees, unless otherwise noted. The impact of recombination in the phylogenetic trees was assessed using ClonalFrameML v.1.12, with default settings.^37^ Trees were visualized and annotated with metadata in FigTree v.1.4.4 and/or iTOL.^38^ Amino acid sequences of RtxA, RtxB, RtxC, RtxD, and TolC from *K. kingae* KWG-1 were used to query the BlastP database on NCBI. Representative sequences of the closest homologs from Gram-negative bacteria were downloaded and aligned using MUSCLE in the MEGAX: Molecular Evolutionary Genetics Analysis version 10 software suite.^39^ Aligned sequences were used for phylogenetic reconstruction using RAxML with 100 bootstraps, the protein GTR substitution model, and the Jones-Taylor-Thornton model. Trees were visualized using iTOL.

### Genotyping clinical isolates

Primers used in this study are listed in Table 1. *K. kingae* strain KK01 (a non-spreading, non-corroding stable derivative of clinical isolate 269-492) was used to design primers against the four ends of the *rtx* loci and the previously described capsule typing primers.^40^ PCR assays were performed on gDNA extracted as described above. For the *rtx* loci, the presence of an amplicon of the expected size on an agarose gel was used to determine presence or absence. The multiplex capsule typing assay produces amplicons of different size for each of the four capsule types.^40^ A gene deletion mutant was used as a negative control to demonstrate PCR specificity. Additional strains not included in sequencing but genotyped by PCR are listed in Table S3. To determine the association between these genotypes and disease state, a Fisher’s exact test was performed in Graphpad Prism v.9.2.0.

**Table 1.**
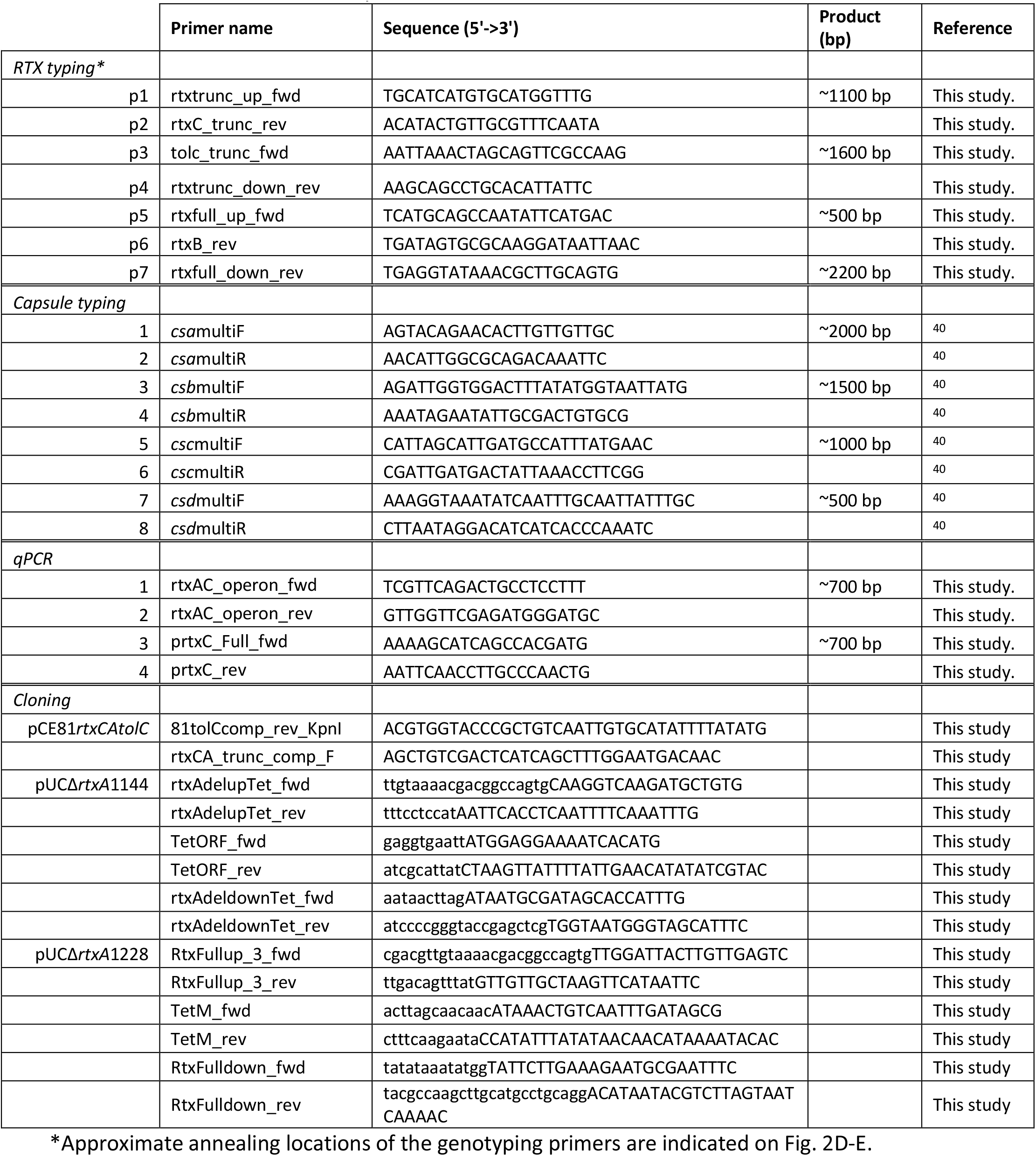
Primers Used in this study.

### *In vitro* characterization of clinical isolates

*K. kingae* isolates used in genotyping PCR assays were grown on chocolate agar at 37°C with 5% CO_2_, as previously described.^9^ *K. negevensis* strains were grown on brain heart infusion (BHI) agar supplemented with 10% sheep blood for 24 hours at 37°C, 5% CO_2_. ^41^

Mutants were generated as described previously.^9,12,42^ Briefly, NEB Builder was used to design primers for three fragment Gibson assembly cloning (Table 2). Plasmids were constructed in the pUC19 backbone using NEBuilder Hi-Fi DNA Assembly Mastermix (New England Biolabs) following the manufacturer’s instructions. Vectors were linearized and subsequently introduced into *K. kingae* by natural transformation followed by selection on chocolate agar plates supplemented with the appropriate antibiotic. Mutations were confirmed by preparing gDNA, amplification by PCR, and Sanger sequencing.

**Table 2.**
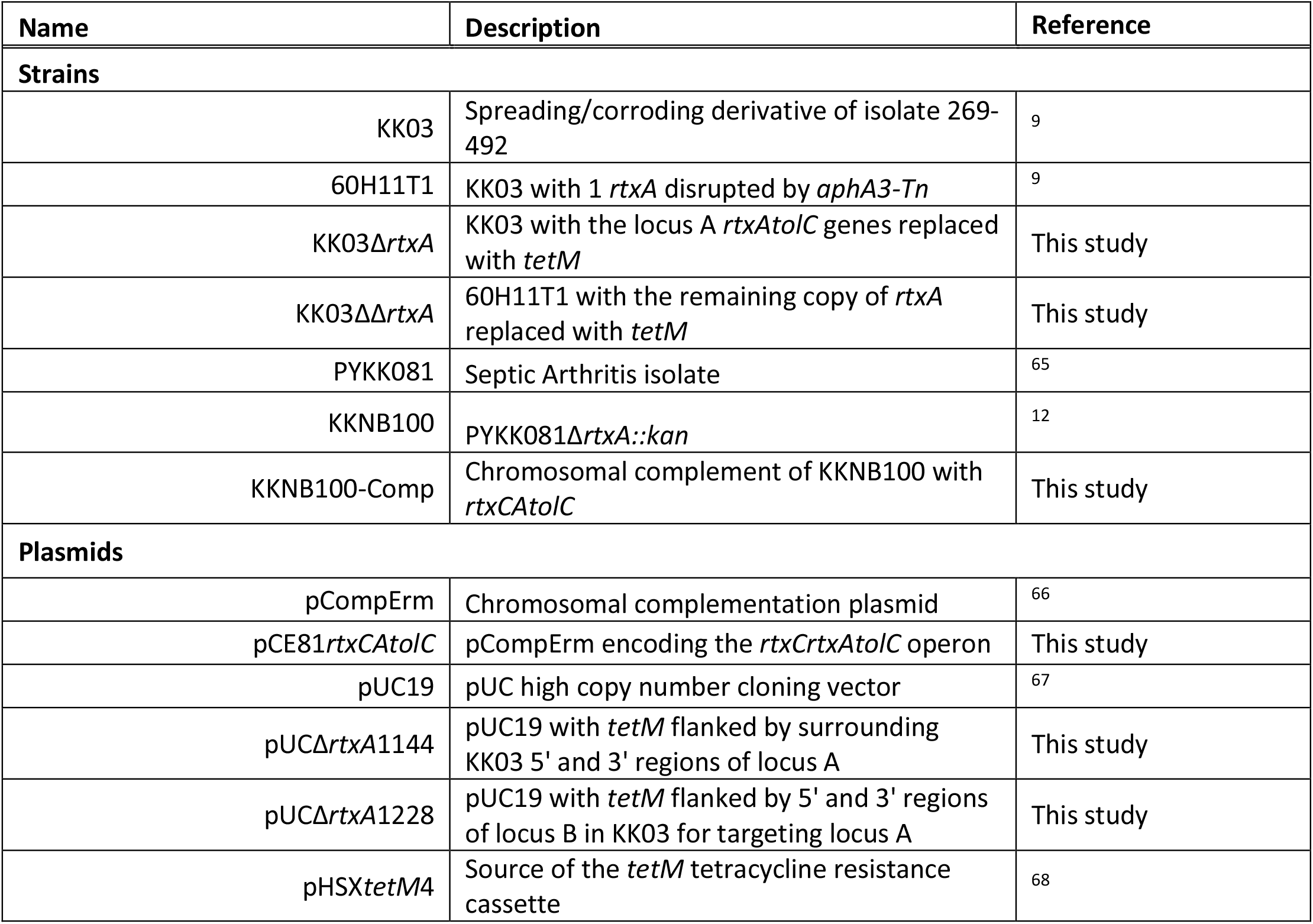
Strains and constructs used in this study.

To complement *rtxA* in PYKK081, the primers “81tolCcomp_rev_KpnI” and “rtxCA_trunc_comp_F” were used to amplify *rtxCrtxAtolC*, and the resulting amplicon was digested with KpnI/SalI and ligated into KpnI/SalI-digested pCompErm, generating pCE-81*rtxCAtolC* (Tables 1-2). The complementation plasmid was linearized with SfoI and transformed into KKNB100 via natural transformation. To generate KK03ΔΔ*rtxA* and KK03Δ*rtxA, rtxA*_LocusA_ was replaced with the *tetM* tetracycline resistance cassette using a four-fragment assembly and the NEB Hifi Assembly Kit. The resulting plasmid was linearized and transformed into an existing *rtxA* transposon mutant (60H11T1) to generate KK03ΔΔ*rtxA* or into strain KK03 to generate KK03Δ*rtxA*.

### RNA extraction

Bacteria were grown for 20 hours at 37°C with 5% CO_2_, resuspended in 3 mL brain heart infusion (BHI) broth to an OD_600_ of 0.8, and incubated at 37°C with 5% CO_2_ for 1 hour. A volume of 1 mL was removed and centrifuged for 15 min at 4,000 x*g* to pellet bacteria. Pellets were resuspended in 1 mL of TRIreagent and incubated for 10 minutes at room temperature (RT). RNA was extracted from the aqueous phase by the addition of 200 μL of chloroform and centrifugation at 10,000 x*g* for 10 minutes. The aqueous phase was removed and added to 600 μL of 100% isopropyl alcohol. Glycogen (ThermoFisher) was added to each sample to aid in pellet visualization, and samples were incubated for 10 minutes at RT. Nucleic acids were pelleted by centrifugation at 10,000 x*g* for 30 minutes. The pellets were washed with 1 mL of 70% ethanol prepared with DEPC-treated water and were then treated with DNAse I to remove contaminating DNA. DNase-treated RNA was then re-extracted as described above. cDNA was generated using qScript cDNA Supermix (Quanta Bioscience) and 1 μg of RNA. Quantitative PCR (qPCR) was performed using primers listed in Table 1.

### Polyclonal antiserum generation

Purified recombinant RtxA (a gift from Nataliya Balashova) was sent to Cocalico Biologicals, Inc. (Stevens, PA) for polyclonal antiserum generation in a guinea pig according to their standard antibody production protocol. The resulting RtxA-specific polyclonal antiserum is designated GP-23.

### Western blotting

*K. kingae* strains were resuspended into heart infusion (HI) broth to an OD_600_ of 0.8 and incubated at 37°C with 5% CO_2_ for 1 hour. A 1 mL aliquot was removed and centrifuged at 14,000 x*g* for 2 minutes. To the clarified supernatant, trichloroacetic acid (TCA) was added to final concentration of 10%. After 15 min on ice, the precipitated proteins were collected by centrifugation at 21,000 x*g* for 10 min at 4°C and were washed with 1 ml ice-cold acetone. After centrifugation at 21,000 x*g*, the resulting pellet was resuspended in 1 x SDS-PAGE loading buffer and boiled for 5 min, and an aliquot was separated using 10% SDS-PAGE. After transfer to nitrocellulose, the blot was blocked in 5% skim milk in 1 x PBS. The blot was then probed with a 1:2,000 dilution of GP23 for 1 hr at ambient temperature, washed with Tris-buffered saline supplemented with 0.1% Tween-20, probed with a 1:10,000 dilution of an anti-guinea pig-horseradish peroxidase-conjugated secondary antibody (Sigma; product no. A5545) for 30 min at ambient temperature, washed again, and developed using a Syngene G-Box XT4 system (Frederick, MD) after exposure to a chemiluminescent substrate (ThermoFisher Supersignal Plus).

### Hemolysis

Hemolysis was assessed with bacteria grown for 20 hours at 37°C with 5% CO_2_ and resuspended in 3 mL of HI broth to on OD_600_ ∼0.8. Suspensions were then diluted 1:100 in HI and mixed in equal volumes with 1% washed sheep red blood cells in 1x PBS supplemented with 0.492 mM CaCl_2_ and 0.900 mM MgCl_2_. After incubation for 30 minutes at 37°C with 5% CO_2_, each reaction was pelleted at 1000 x*g* for 2 minutes. The absorbance at 415 nm was measured for each supernatant. Cultures were standardized to a maximum hemolysis condition generated by adding water in place of bacteria. Percent maximum hemolysis was calculated for each strain relative to a water lysis control. Statistical analysis was performed in Graphpad Prism v.9.2.0.

### Cytotoxicity

Cytotoxicity was assessed by LDH release from 16HBE-14o-cells. Cells were seeded in 1x MEM supplemented with 10% FBS (Corning) at a density of 2.5×10^4^ in a 96-well tissue culture plates (Greiner) in triplicate and grown at 37°C with 5% CO_2_ for 48 hours. Bacteria were grown for 20 hours at 37°C with 5% CO_2_ and resuspended in 3 mL of HI broth to on OD_600_ ∼0.8. After resuspension, the culture media for the 16HBE-140-cells was replaced with 100μL of prewarmed, serum free 1x MEM. A 5 μL aliquot of the resuspended culture was added (M.O.I. ∼100), and the plates were centrifuged for 2 minutes at 1,000 *xg* followed by incubation for 1 hour at 37°C with 5% CO_2._ The infected cells were once again centrifuged for 2 minutes at 1,000 *xg*, and 50 μL of supernatant was removed for quantification. LDH in the cell supernatants was quantified using the Cytotoxicity Detection kit (Roche), following manufacturer’s instructions. Plates were developed for 40 minutes in the dark, and the absorbance at 490 nm was quantified on a plate reader (PerkinElmer). Percent maximum LDH release was calculated for each strain relative to a 1% Triton X-100 lysis control. Statistical analysis was performed in Graphpad Prism v.9.2.0

### Epithelial Barrier Breach

16HBE-14o-cells were maintained in 1x MEM +10% FBS. To establish semi-differentiated cultures, 2 x 10^5^ cells were seeded onto 33 mm ThinCert Cell Culture Inserts with 8 μm pores (Greiner). Prior to seeding, inserts were coated with an extracellular matrix component mixture comprised of 10 μg/mL human fibronectin (Sigma), 100 μg/mL bovine serum albumin (Fisher), 30 μg/mL PureCol collagen (Sigma) in 1x MEM for two hours at room temperature. Cultures were maintained with apical and basolateral media for 24 hours, at which point the apical media was removed and 400 μL of MEM + 10% FBS was added to the basolateral chamber. Cells were allowed to differentiate at the air-liquid interface (ALI) for an additional 5 days, until reaching a transepithelial electrical resistance of ∼500 Ω. The culture media was then replaced with 1x MEM in both the apical and basolateral chambers. ALI cultures were infected with 10 μL of bacteria (5 x 10^6^ colony forming units (CFUs), multiplicity of infection of 10 after growth for 20 hours at 37°C with 5% CO_2_ and resuspended in HI). An additional 290 μL of 1x MEM was added to the apical chamber to accommodate a probe to monitor transepithelial resistance. At 30 min, 1 hr, and 2 hr post-infection, the basolateral media was collected and plated for CFUs. Statistical analysis was performed in Graphpad Prism v.9.2.0 and significant differences are shown.

### Juvenile rat infection model

*K. kingae* strains were swabbed from chocolate agar plates and resuspended in PBS to a final density of 5.0 x 10^8^ CFU/ml. At least eight, five-day-old Sprague-Dawley rat pups (Charles River Laboratories, Wilmington, MA) were injected via the intraperitoneal route with 0.1 ml of the bacterial resuspensions or PBS alone with a 27 1/2-gauge needle and then returned to their cage with a lactating dam. The experimental groups were housed separately and were monitored for signs of morbidity and mortality twice daily for a total of 5 days. Animals found to be moribund were euthanized by using CO_2_ inhalation followed by secondary decapitation. A Kaplan-Meier curve was constructed, and statistical analyses were performed in Graphpad Prism v.9.2.0.

## Results

### A common ancestor of pathogenic *Kingella* species acquired the RTX toxin

Consistent with existing literature, when the five *Kingella* species were assessed for β-hemolysis on BHI plates supplemented with 10% sheep blood, only *K. kingae* and *K. negevensis* were hemolytic (Fig. 1A), suggesting that the toxin gene was likely in the common ancestor of *K. kingae* and *K. negevensis*, the pathogenic *Kingella* species. Given the lack of *rtxA* in the commensal *Kingella* species, several investigators have suggested that *rtxA* was acquired horizontally by *K. kingae*.^2,9,10^ To better understand the acquisition of the RtxA toxin in *K. kingae* and *K. negevensis*, we first need to understand the genetic history of the toxin and associated machinery in the two pathogenic *Kingella* species.

**Fig. 1.**
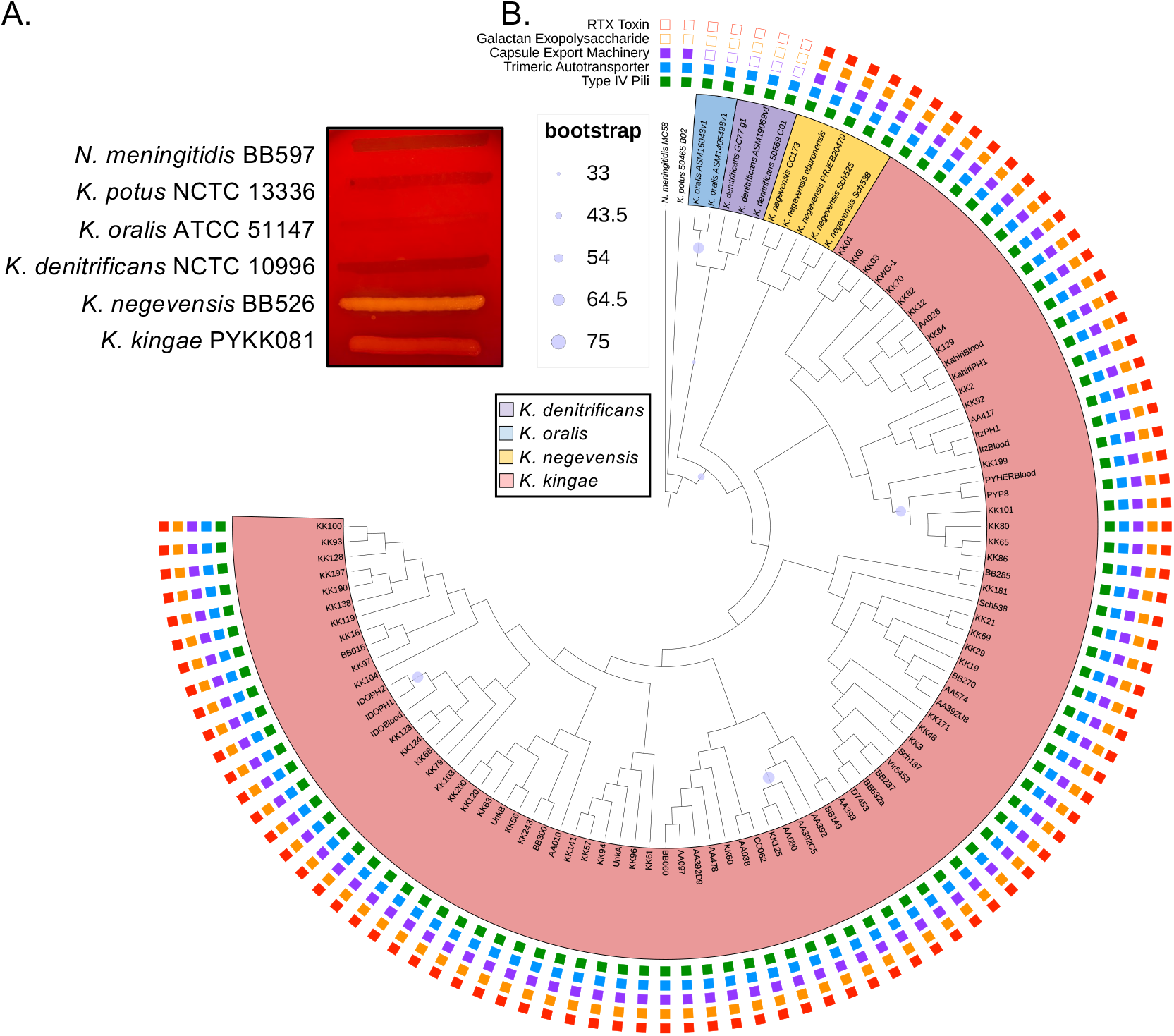
Phylogenetic reconstruction of *Kingella* species. A. Representative strains of *N. meningitidis, K. potus, K. oralis, K. denitrificans, K. negevensis*, and *K. kingae* were grown on blood plates. Only *K. negevensis* and *K. kingae* produce any zone of hemolysis. B. Maximum likelihood phylogeny of all publicly available genomes was reconstructed from whole genome sequences using all *K. kingae* isolates in this study. The phylogeny is rooted by *N. meningitidis* MC58, and the branches and clades are colored by species. Bootstraps N = 100, and bootstrap values less than 75 are indicated by blue circles along the branches. The outer rings denote the presence/absence of each virulence factor utilized by *K. kingae* to facilitate pathogenicity. Closed squares indicate presence of the system, while open squares indicate absence.

To determine the relatedness of *Kingella* strains, a whole genome phylogenetic analysis was performed (Fig. 1B). As expected, *K. negevensis* and *K. kingae* are sister taxa that constitute a single clade of pathogenic *Kingella* species, reinforcing the proposed acquisition of *rtxA* in the common ancestor of these species. Recent work from our group showed that *K. kingae* and *K. negevensis* diverged from *Alysiella* and *Simonsiella* more recently than they diverged from *K. denitrificans*.^43^ Genome analyses showed no evidence of *rtxC* or *rtxA* genes in any of these closely related taxa.

### *K. negevensis* isolates encode the RTX machinery at a single locus

*K. kingae* isolates contain either one or two copies of *rtxCrtxAtolC*.^7,10,13^ To date, little is known about the genomics of *K. negevensis* due to both a lack of available isolates and a lack of whole genome sequencing.^41^ To determine how many RTX loci are present in *K. negevensis*, we performed Oxford Nanopore Technologies (ONT) MinION sequencing on a collection of 12 isolates (Table S1). After sequencing and removal of short reads, we achieved an average read length of ∼13 kB. We performed *de novo* assembly of each sequenced strain, resulting in completed circular genomes and plasmids. Assembled genomes were compared to each of the *K. kingae rtx* genes and complete *rtx* loci using BLAST. We found only a single copy of each gene, with the genes appearing to be syntenic in *K. negevensis*. These data suggest that the genomes of *K. kingae* and *K. negevensis* underwent unique structural changes since diverging. To confirm this conclusion, pooled *K. negevensis* reads were then mapped to *K. kingae* strain KWG-1, which contains *rtx* genes at two loci: locus A (*rtxBrtxDrtxCrtxAtolC*) and locus B (*rtxCrtxAtolC*) (Fig. 2A). We hypothesized that, if the *K. negevensis rtx* system is organized like it is in *K. kingae* strains that contain a single copy of *rtxA*, then *K. negevensis* isolates with a single copy of *rtxA* would show no read through from *rtxD* to *rtxA*, as these genes are non-syntenic in single *rtxA* copy *K. kingae* isolates. After mapping reads, we identified a large population of reads that span from *rtxD* to *rtxA* and no increase in read depth in the duplicated genes (Fig. 2B-C). These data suggest that there is a single copy of *rtxA* in the sequenced *K. negevensis* isolates and that the RTX machinery is encoded by a single locus.

**Fig. 2.**
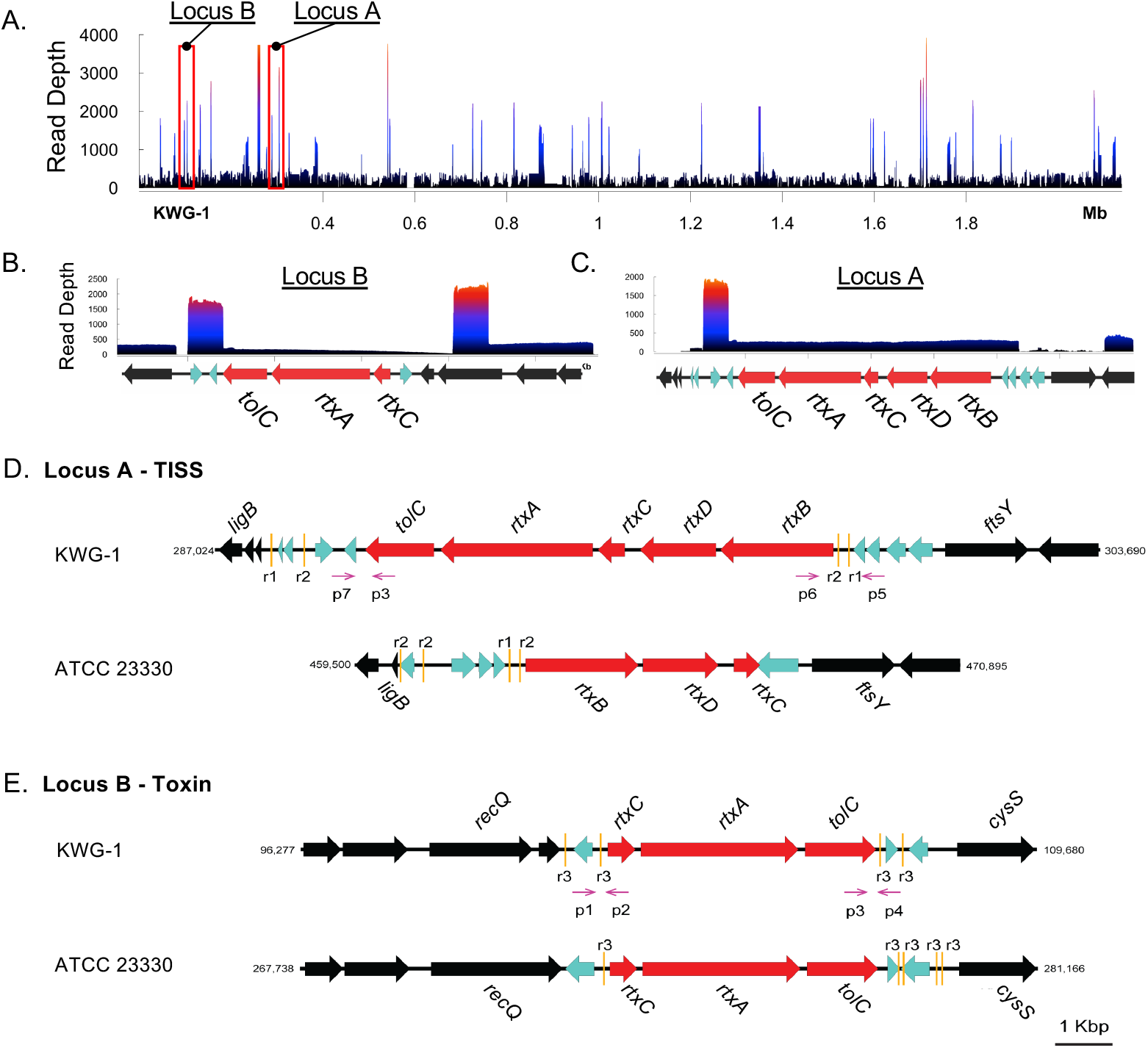
*K. negevensis* encodes the RTX toxin machinery at a single locus. A. Reads from MinION long run sequencing of 12 *K. negevensis* isolates were pooled, mapped to the KWG-1 genome, and visualized using the Sushi package in R. The two RTX toxin loci are highlighted in the red boxes. B-C. Read depth of *K. negevensis* sequencing mapped to locus B and locus A of KWG-1, respectively. The genes found in each region of interest of the KWG-1 genome are diagrammed below the plots. D-E. Genomic maps of the *rtx* loci of representative *K. kingae* genotype I and II isolates (ATCC23330 and KWG-1, respectively). RTX-associated genes are shown in red. Each of these regions is flanked by several putative DNA transposition genes, shown in teal. Gold bars mark repeat sequences flanking each of these loci. Core genes are shown in black. Genotyping primers anneal at each end of these loci and are denoted by magenta arrows (p1-p7).

### *K. kingae* RTX machinery genes are conserved across two loci in all isolates

In all *K. kingae* isolates, we found the *rtx* genes in two loci, named locus A and locus B (Fig. 2D-E). In some isolates, locus A contains the genes encoding the type I secretion system (TISS; *rtxB* and *rtxD*) followed by *rtxC*, and locus B contains an additional copy of *rtxC*, followed by *rtxA* and *tolC* (Fig. 2D), an arrangement designated here as genotype I (Fig. 2D). Both copies of *rtxC* are preceded by a putative promoter for the *rtxCrtxAtolC* operon. Additionally, sequence analysis suggests that the duplicated genes are identical. Consistent with this finding, we confirmed expression of transcripts that contain primarily *rtxC* and *rtxA in vitro* by RT-PCR (Fig. 3A), strongly supporting operonic transcription of these genes, independent of the TISS. In other isolates, locus A contains an additional copy of *rtxA* and *tolC*, an arrangement designated genotype II and described previously in *K. kingae* strain KWG-1 (Fig. 2E).^10,44^

**Fig. 3.**
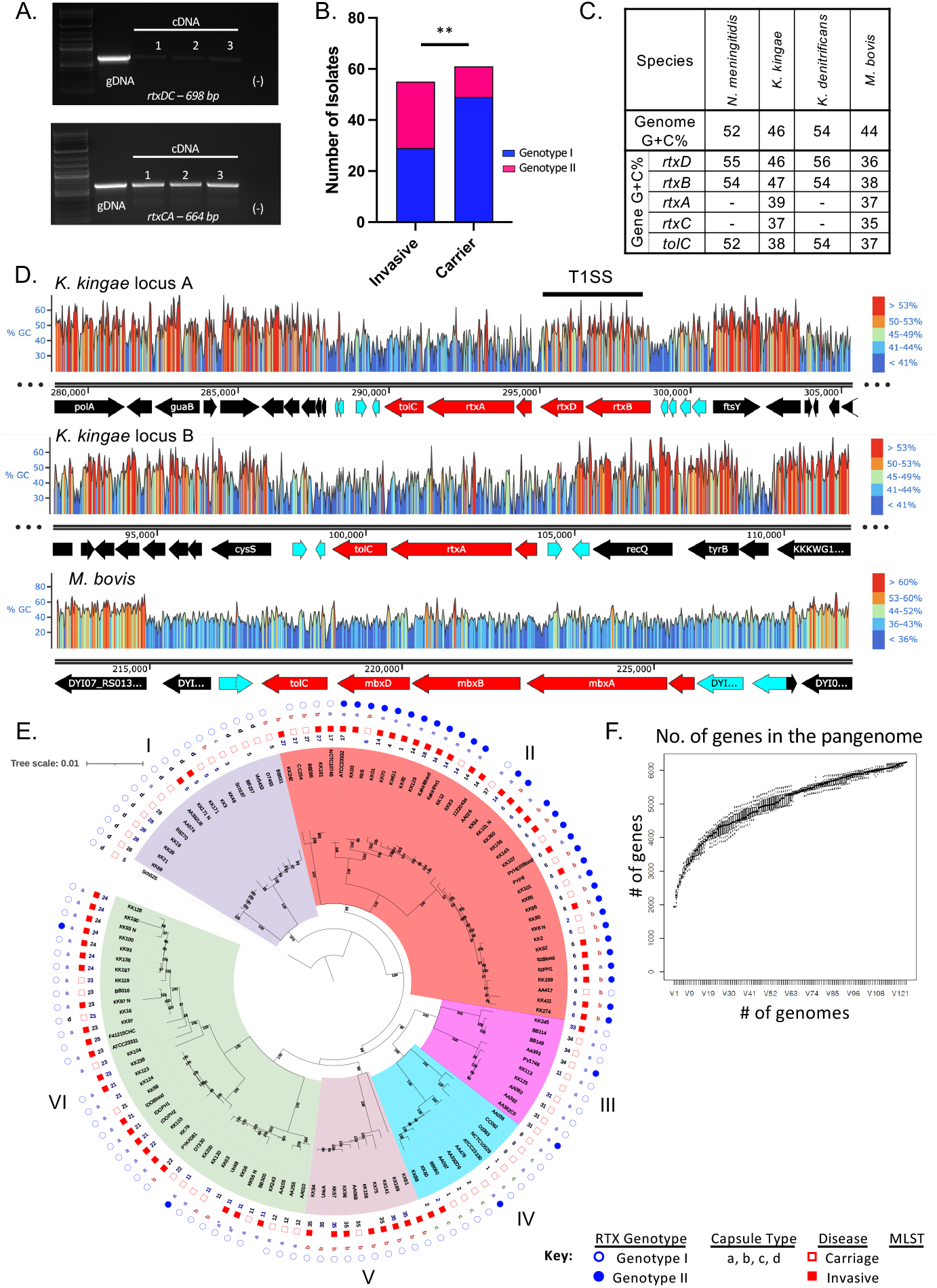
*Reconstruction of the RTX machinery by K. kingae.* A. To determine if these genes are transcribed as an operon in KWG-1, RNA was extracted, and RT-PCR was performed to identify read-through between the *rtxD* and *rtxC* genes (top) and between *rtxC* and *rtxA* (bottom). As a positive control, gDNA extracted from KWG-1 is shown. B. Presence of genotype I or II in a panel of 124 clinical isolates was determined using PCR screening approach. Approximately 30% of the screened isolates are genotype II. Isolates were then stratified by isolation condition, invasive or carriage, and a *Χ*^2^ test of association was performed. There is a significant association between *rtxA* copy number and disease state at the time of isolation (*p* = 0.02, *df = 4*). C. Gene G+C% for each gene associated with the production of RTX family proteins from diverse bacterial species. G+C% for *rtxC* and *rtxA* are only shown for those species in which a close homolog could be identified. D. Snapgene was used to generate a heatmap of G+C% across the toxin associated genes in *K. kingae* strain KWG-1 locus A (top), KWG-1 locus B (middle), and the *M. bovis mbx* pathogenicity island (bottom). Genome maps are labeled below the heatmap, with toxin genes colored red, DNA transposition genes colored cyan, and core genes colored in black. E. A whole genome phylogeny of *K. kingae*, rooted by *K. negevensis* Sch525, was constructed using publicly available genomes and an additional 88 genomes sequenced for this study. For isolates that appear in both the in-house and public databases, leaves marked with a “_N” denote those retrieved from NCBI. Each strain is marked with sequence type (ST), isolation condition (invasive or carriage), capsular polysaccharide type, and *rtx* genotype, as denoted in the key. The phylogeny has six distinct clades (named I-VI), which contain specific ST and polysaccharide capsule types. F. A sampler’s curve of unique genes identified in the pangenome of all *K. kingae* isolates included in this study is shown. The curve quickly saturates and begins to plateau, suggesting that this strain collection represents much of the genomic diversity of clinical *K. kingae* isolates.

To determine the prevalence of the genotype I and genotype II arrangements, we designed a PCR-based approach that would specifically amplify the 5’ end of the additional copy of *tolC* found in locus A of genotype II isolates (Fig. 2D-E). A total of 124 clinical *K. kingae* isolates that were collected primarily in Israel between 1990 and 2012 were analyzed (Table S1). Approximately 30% of these isolates contain genotype II *rtx* loci, suggesting that genotype II is prevalent in the population. When strains were stratified by site of isolation, genotype II was strongly associated with strains recovered from patients with invasive disease (Χ^2^ test of association; *p*=0.02; *df*=2) (Fig. 3B). A second screen was performed using an additional 131 global isolates collected between 1986 and 2015, revealing genotype II in approximately 35% of these isolates (Table S3).

### Acquisition and reconstitution of fully functional RTX machinery

The *rtx* loci in *K. kingae* have features consistent with mobile genetic elements. Both loci are flanked by at least one putative transposon gene and repetitive sequences, and both have a G+C% that is lower than the G+C% in the rest of the genome (Fig. 2D-E, 3C-D). The putative transposon genes belong to the IS5 family of transposases (Fig. 2D-E).

To better understand the horizontal gene transfer event that resulted in the acquisition of *rtxA*, we performed a phylogenetic analysis of the five proteins involved in toxin production and secretion, including the RtxA toxin, the RtxC toxin-activating acyltransferase, and the RtxB, RtxD, and TolC components of the TISS. The NCBI RefSeq database was queried using the predicted amino acid sequences of each protein in *K. kingae* strain KWG-1. Homologous sequences were downloaded and used for phylogenetic reconstruction, allowing identification of RTX-associated proteins across 14 diverse bacterial genera (Fig. S1). The reconstructions of the RtxA, RtxC, and TolC phylogenies are largely congruent and demonstrate that these *K. kingae* proteins are most closely related to homologs in distantly related *Moraxella* species (Fig. S1A-B, E).

Importantly, there are no close homologs to RtxA or RtxC in other species in the *Neisseriaceae*. Instead, the closest orthologous sequences are the Frp proteins in *N. meningitidis* and predicted Frp-related gene fragments in the non-pathogenic *Neisseria* species (Fig. S1A-B). This pattern contrasts starkly with the RtxB and RtxD phylogenies, where the closest homologs to the *K. kingae* proteins are in the *Neisseriaceae* and include homologs in several other *Kingella* species and *Alysiella crassa* (Fig. S1C-D). A full list of protein identities and similarities for proposed homologs in *K. kingae, K. oralis, K. denitrificans, K. negevensis, M. bovis*, and *N. meningitidis* is shown in Table S4. In addition, the G+C% content of the *rtxD* and *rtxB* genes in *K. kingae* (46% and 47%, respectively) is similar to the average G+C% of the *K. kingae* genome (46% for KWG-1) (Fig. 3C). The G+C% content of the *rtxC, rtxA*, and *tolC* genes at both loci in KWG-1 is considerably lower (Fig. 3C-D), consistent with the putative acquisition of these genes through horizontal gene transfer. Together these findings strongly suggest that the *rtxA, rtxC*, and *tolC* genes have an independent origin from *rtxB* and *rtxD* and that the full complement of genes was reconstituted as a functional toxin production system in the common ancestor of the pathogenic *Kingella* species.

### Whole genome sequencing of *K. kingae* isolates

To investigate the role of the toxin duplication in *K. kingae* and to test for other factors associated with invasive disease, we sequenced 74 *K. kingae* and two *K. negevensis* isolates with Illumina HiSeq and combined these sequences with the genomes of 52 *K. kingae* strains in the NCBI Sequence Read Archive (SRA) (Table S1). The average genome size for *K. kingae* is 2,003,405 bp (range: KK120, 1,943,140 bp; PYP8, 2,116,086). The average G+C% for the analyzed *K. kingae* strains is 46.75% (range: NCTC10746, 46.39%; KK242, 46.90%, assembly statistics data are listed in Tables S2).

As shown in Fig. 3E, a whole-genome phylogeny of *K. kingae* strains was constructed. For each isolate, multilocus sequence type (MLST), polysaccharide capsule type, and clinical condition of the patient was mapped to the tree. The resulting tree can be divided into six clades (I-VI). The clades diverge along MLST and between capsule types. Clade I is comprised of primarily capsule type d carriage isolates. Clades IIa, VI, and VII contain primarily capsule type a isolates recovered from patients with invasive *K. kingae* disease. Clades IIb and V contain primarily capsule type b isolates from patients with invasive disease. Clades III and IV contain primarily capsule types a and c carriage isolates. Isolates do not group by isolation location or year. Of the two clades that are enriched for invasive isolates, clade II is also enriched for *rtx* genotype II isolates, raising two possibilities: (1) the duplication of the *rtxA* gene is important for facilitating *K. kingae* pathogenesis, or (2) there is another genomic factor enriched in this clade that enhances invasion.

### Pangenome analysis reveals a limited pangenome and no other genes clearly associated with pathogenicity

To determine if there are additional genes that enhance virulence in isolates from invasive clades, we analyzed the pangenome using the Roary pipeline. A total of 6241 gene clusters were identified across these 124 isolates (Tables S5-S6). Interestingly, though *K. kingae* is naturally competent, in a collectors’ curve analysis, each new genome beyond 75 added very few new genes to the pangenome, suggesting that we have covered much of the pangenomic diversity of this species (Fig. 3F). The core genome is comprised of 1188 genes found in more than 95% of isolates (Table S5). The pangenome is comprised of 5053 genes, including 3720 that are found in less than 15% of genomes. To determine which genes may be associated with invasive isolates, Scoary was used to perform a genome wide association study comparing invasive and carriage isolates, and comparing between clades (Table 3, S7). Eleven genes are specifically enriched in invasive isolates. As few of these genes have been characterized in *K. kingae*, functional predictions were made using BLASTp and HHPRED (Table 3), and none of these genes has a clear role in pathogenicity.

**Table 3.** Pangenome genes associated with Invasive and Carriage Isolates.

|  | *Gene* | *Odds_ratio* | *Annotation* |
| --- | --- | --- | --- |
| **Enriched in Invasive** | KKKWG1_0747 | 0.13919414 | Inovirus associated |
|  | KKKWG1_1631 | 0.16053512 | *hp* |
|  | KKKWG1_1448 | 0.17085714 | Peptidoglycan transpeptidase |
|  | KKKWG1_0749 | 0.19285309 | Inovirus associated |
|  | KKKWG1_0182 | 0.17769608 | Transpeptidase (NTF2-like) |
|  | ABJGGOKB_01117 | 0.13425254 | Pentapeptide repeat |
|  | KKKWG1_0242 | 0.15384615 | Ribonuclease VapC |
|  | KKKWG1_0241 | 0.15384615 | *hp* |
|  | KKKWG1_0029 | 0.19962453 | ABC Transporter |
|  | ABJGGOKB_00731 | 0.12882448 | Lytic Transglycosylase SLT domain |
|  | PIGKNEIA_01059 | 0.21125 | Inovirus associated |
| **Enriched in Carriage** | ABJGGOKB_00216 | 12.88288288 | Transposase |
|  | ABJGGOKB_00217 | 12.88288288 | Endonuclease NucS |
|  | ABJGGOKB_00814 | 24.61363636 | Restriction Endonuclease |
|  | ABJGGOKB_01644 | 24.61363636 | Transposase |
|  | ABJGGOKB_01645 | 22.8 | Acyl-carrier protein |
|  | ABJGGOKB_01643 | inf | *hp* |
|  | ABJGGOKB_00501 | 13.02325581 | DNA binding cold-shock protein |
|  | ABJGGOKB_00500 | 13.02325581 | DMT family |
|  | ABJGGOKB_01824 | 13.02325581 | Helix-turn-helix transcriptional regulator |
|  | ABJGGOKB_01823 | 13.02325581 | Endopeptidase |
|  | ABJGGOKB_01822 | 13.02325581 | *hp* |
|  | KKKWG1_1450 | 9.837398374 | *hp* |
|  | ABJGGOKB_01646 | inf | *hp* |
|  | ABJGGOKB_00278 | 8.307692308 | *hp* |
|  | ABJGGOKB_00732 | 9.166666667 | Lytic Transglycosylase LysM |
|  | JIMKOJHD_01738 | 7.243902439 | GntP Family permease |

Note: "*hp*" denotes hypothetical proteins with no predicted homologs in the BLAST database.

### *rtxA* copy number in isogenic strains influences toxicity in vitro but not virulence in animals

The RtxA toxin is required for virulence in an animal model of invasive disease using strain PYKK081, a genotype I isolate.^10,12^ To examine whether duplication of *rtxA* influences virulence, we compared strains PYKK081, KKNB100 (PYKK081Δ*rtxA*), KK03 (genotype II), KK03ΔΔ*rtxA*, and KK03Δ*rtxA* (genocopies a genotype I). After a 1-hour incubation at 37°C with 5% CO_2_, we observed high levels of RtxA in culture supernatants of PYKK081 and KK03. In contrast, KKNB100 and KK03ΔΔ*rtxA* secreted no RtxA. Secretion of RtxA was rescued by chromosomal complementation of *rtxCrtxAtolC* in KKNB100. KK03Δ*rtxA* secretes less toxin than KK03 (Fig. 4A). As shown in Fig. 4B, PYKK081 and KK03 are highly hemolytic; hemolysis was eliminated in KKNB100 and KK03ΔΔ*rtxA* and was rescued by complementation of *rtxCrtxAtolC* in KKNB100. Hemolysis by KK03Δ*rtxA* was significantly reduced relative to KK03 (Two-way ANOVA, *p*=0.03). Cytotoxicity was assessed by LDH release from 16HBE-14o-human bronchial epithelial cells and was similar between KK03 and KK03Δ*rtxA* (Fig. 4C), likely due to the high sensitivity of these cells to this toxin *in vitro*.

**Fig. 4.**
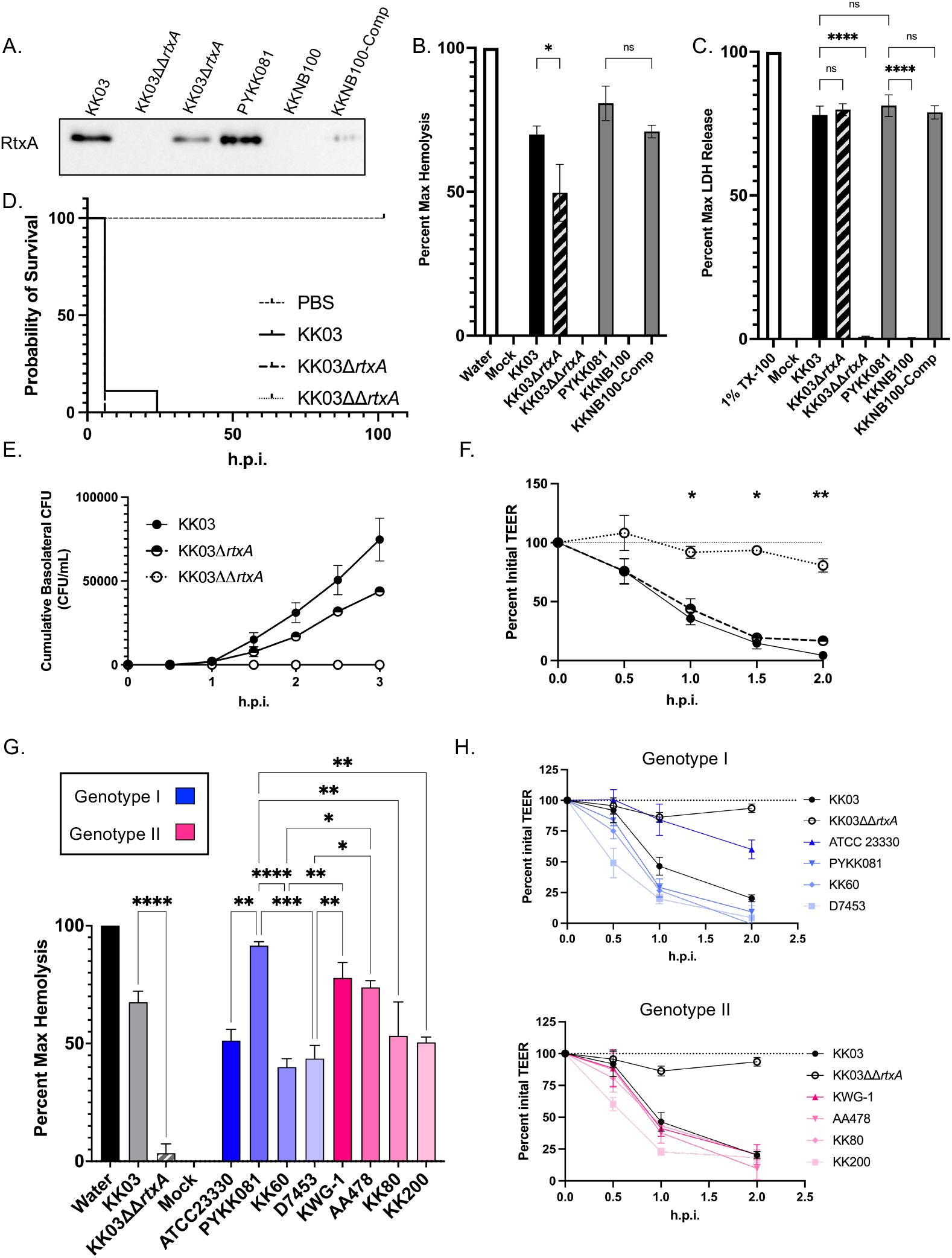
*In vitro and in vivo characterizations of genotype I and genotype II isolates.* A. RtxA levels secreted into the culture supernatant were determined via Western blot. To quantify differences in toxin activity, Liquid hemolysis of sheep red blood cells and C. LDH release from 16HBE14o-cells was examined using isogenic genotype I and genotype II strains. PYKK081, KKNB100, and KKNB100-comp are shown as controls. B-C. Statistics were calculated with a one-way ANOVA \**p*<0.05, \*\**p*<0.01, \*\*\**p*<0.001, \*\*\*\**p*<0.0001. D. Kaplan-Meier survival curve of juvenile rat pups infected i.p. with KK03, KK03Δ*rtxA*, or KK03ΔΔ*rtxA*. PBS and KK03ΔΔrtxA are significantly different from WT (*p*<0.001) and KK03Δ*rtxA* (*p*<0.001). Statistics were calculated by the Mantel-Cox test, N=9 for all groups. E-F. 16HBE14o-cells were cultured at an ALI and infected at a multiplicity of infection of ∼10 on the apical surface in 1x MEM. Over the course of infection, cumulative transited CFUs in the basolateral chamber (E) were monitored every 30 minutes for three hours post infection (h.p.i). Transepithelial resistance (F) was monitored every 30 mins for 2 h.p.i. KK03 is shown in filled circles, KK03Δ*rtxA* is shown in half filled circles, and KK03ΔΔ*rtxA* is shown in open circles. G. To quantify differences in RtxA efficacy, the same panel of clinical isolates was used in liquid hemolysis assays using sheep RBCs. Hemolysis was quantified by calculating the percent relative to water (maximum lysis). Genotype I isolates are shown in blue, and genotype II isolates are shown in pink. Statistics calculated with a one-way ANOVA \**p*<0.05, \*\**p*<0.01, \*\*\**p*<0.001, \*\*\*\**p*<0.0001. H. TEER was monitored over the course of infection of 16HBE-14o-semi-differentiated monolaters for genotype I (top) and genotype II isolates (bottom). Dashed line represents 100% initial TEER. For G-H, KK03 and KK03ΔΔ*rtxA* are shown as positive and negative controls, respectively. All *in vitro* experiments include at least 3, independent biological replicates, averages are plotted with error bars representing the standard error of the mean (±1 SEM).

To assess the impact of these genotypes on invasive disease, we employed the juvenile Sprague Dawley model of disease. Five-day-old rats were inoculated intraperitoneally with 5 x 10^7^ colony forming units of KK03, KΚ03Δ*rtxA*, or KK03ΔΔ*rtxA* and monitored for five days. With toxigenic strains, we observed significant death in the first 24 hours post infection (h.p.i) (Fig. 4D). All animals infected with KK03 or KK03Δ*rtxA* succumbed to infection, while animals infected with KK03ΔΔ*rtxA* or mock infected survived (Mantel-Cox test, both *p*<0.0001). There was no difference in virulence between KK03Δ*rtxA* and KK03 (Mantel-Cox test).

To address whether RtxA influences bacterial invasion across the epithelial barrier, we adapted a model used for other respiratory pathogens to assess *in vitro* epithelial barrier integrity using 16HBE-14o-cells.^45,46^ 16HBE-14o-cells were cultured at an air-liquid interface (ALI) over the course of 10 days, allowing transepithelial resistance (TEER) to steadily increase. Cells were then transferred to serum-free media and infected apically with strain KK03 and the KK03 isogenic mutants and with strain PYKK081 and the PYKK081 isogenic mutant. As shown in Fig. 4E-F and Fig. S2A-B, RtxA-producing strains crossed the epithelial barrier, while strains KK03ΔΔ*rtxA* and KKNB100 were attenuated. Strain KK03Δ*rtxA* transited the epithelial barrier slower than KK03 (Fig. 4E). The strain background had a significant influence on the kinetics of this process, with PYKK081 crossing much faster and in higher numbers than KK03. The decrease in TEER correlated with transited CFUs (Fig. 4F, S2B). Both WT strains caused a rapid drop in TEER, indicative of significant disruption of tight junctions and a loss of epithelial barrier integrity. In contrast, strains KK03ΔΔ*rtxA* and KKNB100 caused no significant change in TEER over the course of infection (Fig. 4F).

### Diverse genotype II isolates are not more pathogenic *in vitro*

To further investigate the impact of genotypes I and II on virulence, we selected eight clinical isolates from different clades across the *K. kingae* population structure (Fig. S2C). All of these isolates secreted similar levels of RtxA into the supernatant, suggesting that genotype II isolates do not generally produce more toxin than genotype I isolates (Fig. S2D). To address differences in activity after secretion, hemolysis and cytotoxicity assays were performed for each strain, revealing no consistent differences between genotype I and genotype II strains (Fig. 4G, S2E).

To examine the influence of genotype on the process of invasion, we used the model with polarized 16HBE-14o-cells. Cultures were infected apically at multiplicity of infection of ∼10. As shown in Fig. 4H and Fig. S2G-J, over the course of infection, both genotype I and genotype II isolates were able to efficiently disrupt TEER with similar kinetics, resulting in similar bacterial transit (Two-way ANOVA, Fig. S2G-J).

## Discussion

*K. kingae* is an oropharyngeal commensal that is most common in young children and is increasingly recognized as a pathogen capable of causing severe systemic disease.^1,2^ Recent studies have identified five major virulence determinants that are conserved across this species: type IV pili, a trimeric autotransporter adhesin called Knh, a polysaccharide capsule, an exopolysaccharide, and a broadly active RTX cytotoxin called RtxA.^7^ The lipopolysaccharide-associated exopolysaccharide and the RtxA toxin are found only in pathogenic *Kingella* species.^11,47^ In this study, we reconstructed the evolutionary history of *rtxA* in the pathogenic *Kingella* species based on a large collection of isolates from both invasive disease and oropharyngeal carriage.

In *K. negevensis*, the genes encoding the toxin-associated machinery (*rtxBrtxDrtxCrtxAtolC*) are found in a single locus. In contrast, using a novel PCR-based approach on a collection of 255 clinical isolates, we determined that all *K. kingae* strains contain *rtx* genes at two loci, and 30-35% have a duplication resulting in a second copy of *rtxA* and *tolC* downstream of the TISS genes and the *rtxC* gene (Fig. 2D-E). These results are reminiscent of *Actinobacillus pleuropneumoniae*, a porcine-specific, respiratory pathogen that contains several copies of genes encoding the RTX toxin called ApxI-ApxIV, a major virulence determinant in this species.^48–50^

Our analysis suggests that the *rtxCrtxAtolC* locus was acquired horizontally in the common ancestor of *K. kingae* and *K. negevensis*. We hypothesize that the locus was acquired from a relative of *M. bovis* or from *M. bovis* itself. *M. bovis* is a bovine respiratory tract pathogen and employs an RTX toxin as a major virulence factor,^51,52^ and in toxin-associated genes present in *M. bovis* were also likely acquired as part of a genomic islet in a single event.^53^ While the *M. bovis* islet contains a full toxin production and secretion system, acquisition of the *rtxCrtxAtolC* locus alone would have resulted in a nonfunctional toxin production system in the pathogenic *Kingella* species because it lacked a TISS for export. However, our analyses suggest that the presence of a preexisting TISS (*rtxBrtxD*) allowed reconstitution of a complete toxin production and secretion system through the process of co-option.

The preexisting TISS is most closely related to genes of the *frp* operon, which encodes the distantly related RTX family protein, FrpC, and is intact in *N. meningitidis* but not in *N. gonorrhoeae* or commensal *Neisseria* speices^54,55^. Though the specific function of FrpC is unclear, it is thought to act as an additional adhesin that is dispensable for virulence in a mouse model^56–60^. This important genetic reconstitution event provides a novel example of gene co-option as a major contributing factor in the evolution of bacterial virulence in a commensal genus. Previous work has described co-option of phage-associated proteins for novel functions, including the *Caulobacterales spmX* gene, the type IV and type VI secretion systems, and a bacteriocin secreted by *Pseudomonas syringae*.^61–64^ However, this is the first report that we are aware of which shows co-option of the components required for toxin elaboration. Reconstitution of a complete RTX system by this mechanism is an essential and defining feature in the evolution of invasive disease by this genus.

We sought to test the association between *rtxA* copy-number variability and invasive disease *in vitro* and *in vivo*. Using a panel comprised of four one-copy and four two-copy isolates, we assessed secreted RtxA levels using Western blotting. We then used *in vitro* assays of toxin function, including hemolysis, cytotoxicity, and epithelial barrier breach. Using genotype I and II strains and isogenic derivatives, we observed a reduction in toxin secretion and hemolysis when one copy of the *rtxA* toxin was disrupted. However, we could not differentiate between diverse genotype I and II isolates in a panel of clinical isolates, suggesting that our *in vitro* model may be overly sensitive to other qualities of clinical strains that cloud toxin-specific effects. While our whole genome sequencing suggests that the *rtxA* genes are identical between the two *rtx* loci, the promoters of the proposed *rtxCrtxAtolC* operons are unique to each copy, suggesting that there may be meaningful differences in regulation in two copy isolates under different environmental conditions.

We propose the following model of the acquisition, co-option, and duplication of the *rtx* toxin genes and the associated TISS in *Kingella*. A common ancestor of all genera in the *Neisseriaceae* contained the Frp system and the necessary TISS required for the secretion of the FrpC RTX protein (Fig. 5). Following acquisition, the functionality of the Frp system was maintained only in the lineage that later evolved into *N. meningitidis*. While the TISS remained largely intact, other genera accumulated mutations in FrpC, ablating its activity and resulting in the *frp* fragments that we observe in contemporary bacterial isolates. As such, we identified the components of a homologous TISS in several members of the *Kingella* clade within the *Neisseriaceae*.

**Fig. 5.**
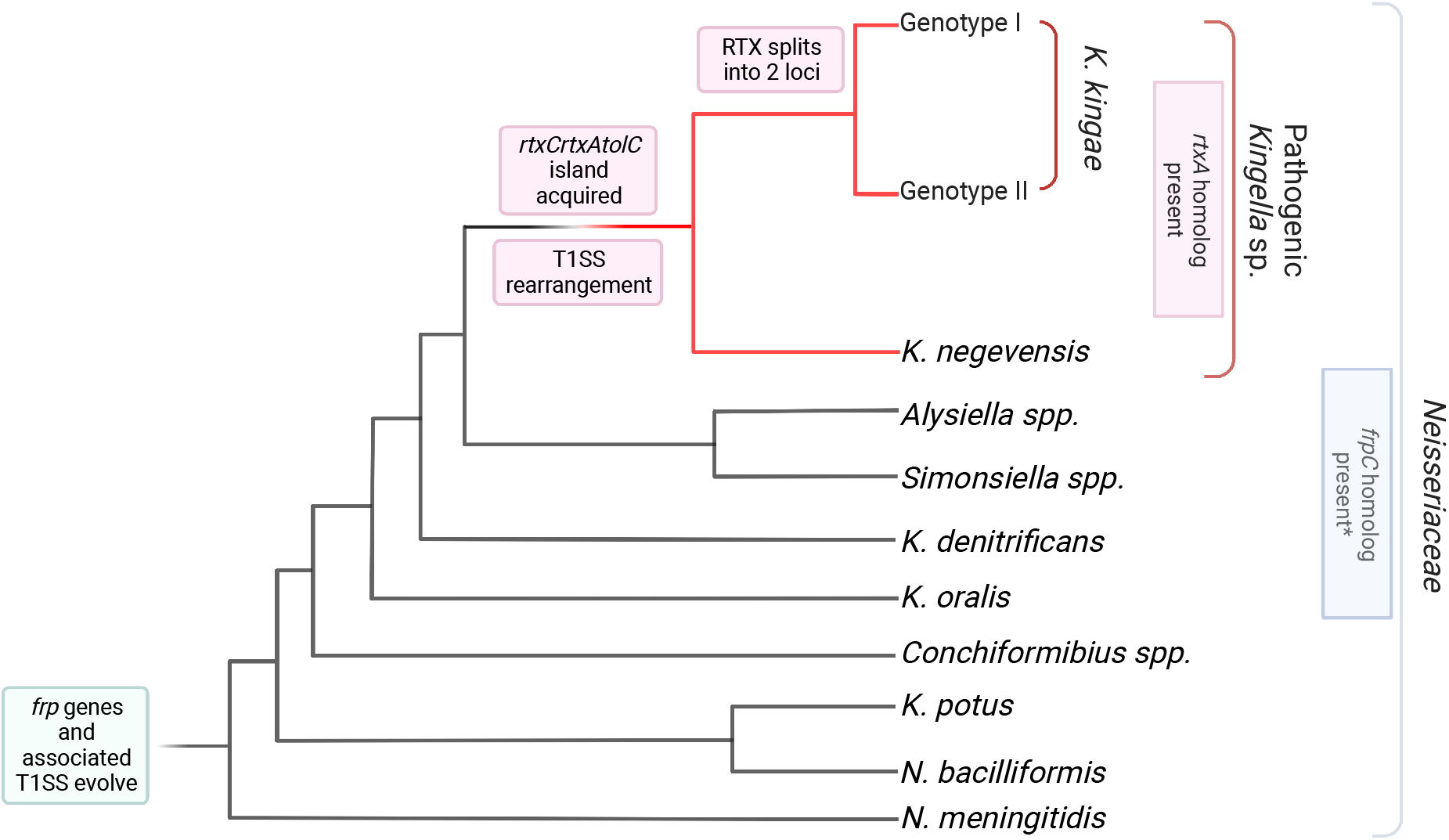
*Model of evolution of the rtx* toxin and the associated T1SS *in K. kingae*. The Frp-associated genes were acquired by the last common ancestor of *N. meningitidis* and *Kingella species* and have been maintained in descendants through the divergence of *N. meningitidis* and the *Kingella* genus. Early branching species of *Kingella* are not associated with invasive disease, though they maintain the Frp machinery. Homologs of the *rtxC* and *rtxA* genes were then acquired by the last common ancestor of *K. negevensis* and *K. kingae*, without an accompanying TISS. The TISS associated with the *frp* genes was co-opted by the *rtxCrtxAtolC* system to both facilitate the development of the pathogenic *Kingella* species and confer a fitness advantage. As *K. kingae* and *K. negevensis* diverge, the RTX machinery underwent two recombination events in *K. kingae* which first split the genes into two loci in the genome, then resulted in the duplication of *rtxC* and *rtxA* in a specific, highly invasive lineage.

The common ancestor of the pathogenic *Kingella* species acquired *rtxCrtxAtolC*. Following acquisition of these genes, we propose that toxin secretion occurred via the existing TISS, which was previously associated with the Frp system. This substantially increased the fitness of this ancestral species and resulted in the appearance of invasive disease by *Kingella* species. Thus *K. kingae* co-opted the FrpC toxin-associated secretion genes, now *rtxBrtxD*, to export RtxA rather than FrpC. Subsequent recombination events brought the TISS into close genetic proximity with the *rtxC* gene, but the *rtxDrtxB* operon remained transcriptionally independent from *rtxCrtxAtolC*, as we observe in *K. kingae*. In *K. kingae*, further reorganization of the genome split the toxin machinery across two distant loci, duplicating the *rtxC* gene. Finally, a clonal group of *K. kingae* associated with invasive disease underwent another round of genomic reorganization, resulting in a second duplication event establishing two copies of *rtxCrtxAtolC* in the genome.

While it is possible that the reconstitution of the RTX machinery occurred in a common ancestor of all *Kingella*, we find it unlikely that the commensal species lost the toxin genes. Additionally, it is possible that *rtxCrtxAtolC* was acquired along with genes for a functional TISS but these genes were subsequently lost, being functionally replaced by the TISS associated with the Frp system. It is interesting to speculate that the secretion system encoded by the *rtxBrtxD* genes may have been conserved over time due to the secretion of other proteins from the bacterial cell, though further work is required to characterize the secretome of *K. kingae* and identify those proteins specifically secreted by the *rtxBrtxD-*encoded TISS.

## Data availability

Files containing phylogenetic trees in Newick format and sequence alignments are available at Figshare (DOI: 10.6084/m9.figshare.21534147). All sequencing data generated in this study is publicly available on the National Center for Biotechnology Information’s (NCBI) Sequence Read Archive (SRA) under Bioproject PRJNA896475. All other data and strains presented in this study are available from the authors by request.

## Ethics statement

The animal procedures were approved by the Children’s Hospital of Philadelphia Institutional Animal Care and Use Committee (IACUC) under protocol IAC 19-001050. The Children’s Hospital of Philadelphia animal research facilities are fully accredited by the Association for Assessment and Accreditation of Laboratory Animal Care (AAALAC) International. The Animal Welfare Act, the Public Health Service policy on the humane care and use of laboratory animals from the U.S. Department of Health and Human Services, and the recommendations in the Guide for the Care and Use of Laboratory Animals of the National Research Council were followed in this work.

## Author Contributions

J.W.S. and P.J.P. secured funding for this work and supervised the study. D.P.M., J.W.S., and P.J.P. conceived of the study. D.P.M., E.A.P., and B.K.K. designed and performed *in vitro* and *in vivo* experiments and analyzed data. D.P.M. conducted *in silico* experiments. D.P.M. E.A.P., J.W.S., and P.J.P. interpreted the data. DPM wrote the original draft of the manuscript. All authors edited and approved the final manuscript.

## Acknowledgements and Funding

We want to thank Dr. Nataliya Balashova for generously providing us with the stain PYKK081Δ*rtxA* (KKNB100) and purified RtxA. We would also like to thank all members of the St. Geme and Planet labs, particularly Dr. Nina Montoya and Kevin Hernandez for their constructive feedback and assistance during the preparation of this manuscript. This work was supported by the National Institute of Allergy and Infectious Diseases under 5T32AI141393 to DPM and 1R01AI121015 to JWS.

## Supplemental Figure Legends

**Fig. S1.**
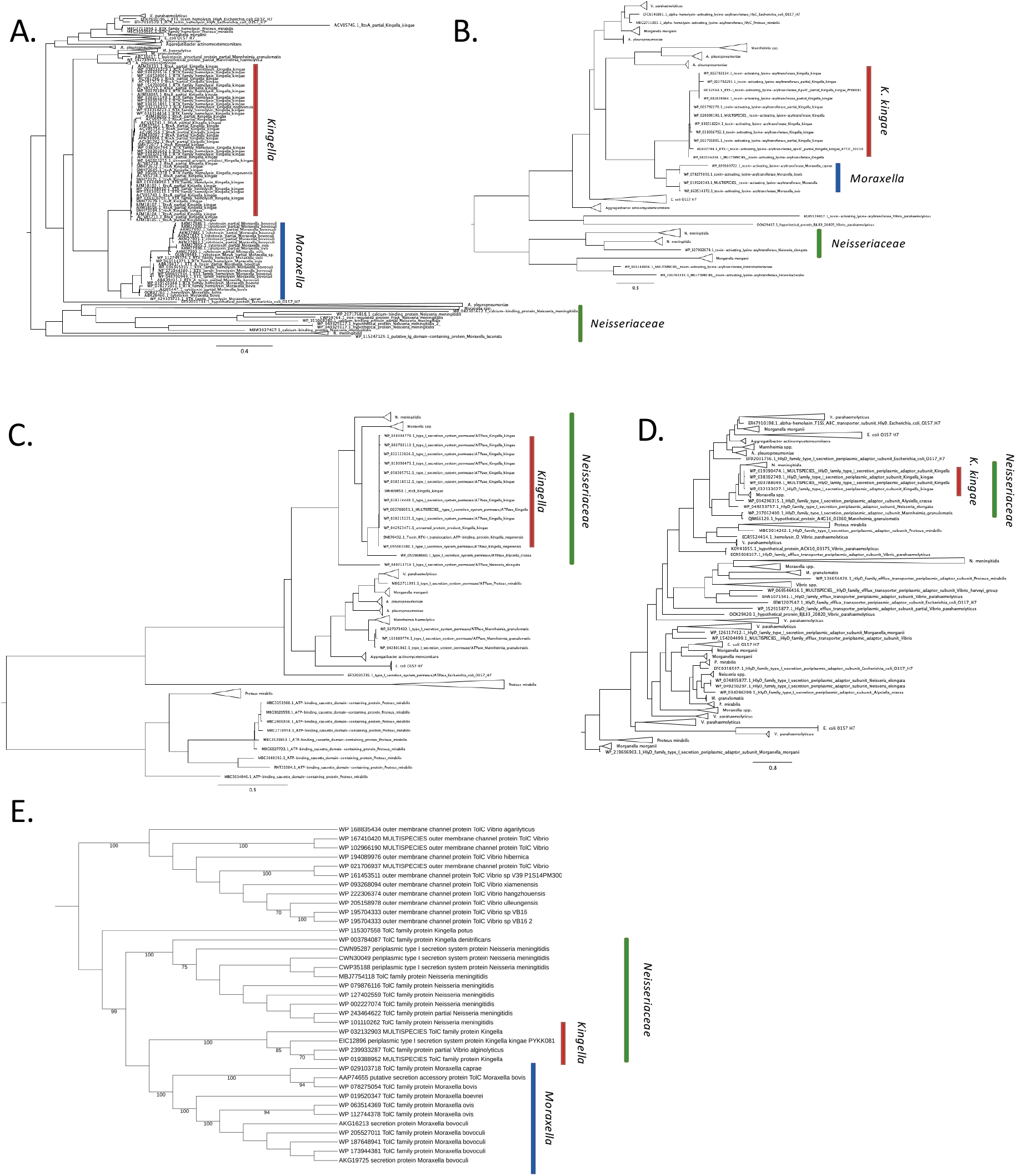
RTX-associated genes show distinct homologs from diverse bacterial species. A-E. BlastP was used to identify homologs of RtxA (A), RtxB (B), RtxC (C), RtxD (D), and TolC (E) in diverse bacterial species. Homologous sequences were downloaded and used for phylogenetic reconstruction allowing us to identify RTX proteins across at least 14 diverse bacterial genera with RaxML. Taxonomic classifications of interest are annotated for each phylogeny. Protein trees are rooted by *E. coli* or *Vibrio* species.

**Fig. S2.**
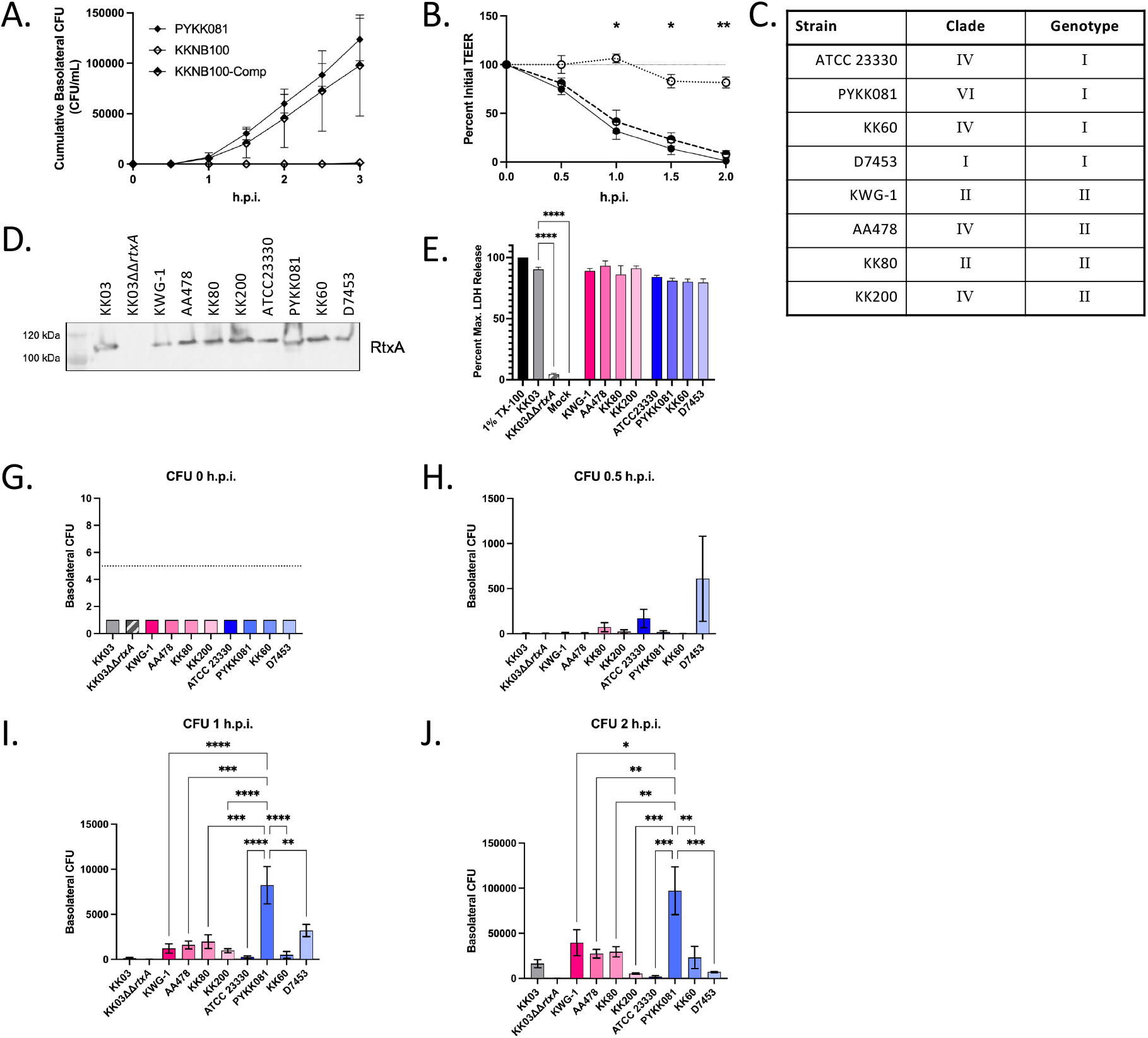
*In vitro* characterizations of RTX copy number variability. A-B. 16HBE14o-cells were cultured at an ALI and infected at a multiplicity of infection (MOI) of ∼10 on the apical surface in 1x MEM with PYKK081, KKNB100 (Δ*rtxA*), or KKNB100-Comp. Over the course of infection, cumulative transited CFUs in the basolateral chamber (A) was monitored every 30 minutes for three hours post infection (h.p.i). Transepithelial resistance (B) was monitored every 30 mins for 2 h.p.i. PYKK081 WT is shown in filled diamonds, KKNB100 (Δ*rtxA*) is shown in open filled diamonds, and KKNB100-Comp is shown in half filled diamonds. C. A clinical panel of isolates was constructed to include four diverse isolates from genotypes I and II, respectively. D. RtxA levels secreted into the culture supernatant were determined via Western blot. E. LDH release by 16HBE-14o-cells was determined using the panel of clinical isolates. Monolayers were infected for 1 hour at an MOI of ∼40. LDH was quantified. Statistics calculated with a one-way ANOVA \**p*<0.05, \*\**p*<0.01, \*\*\**p*<0.001, \*\*\*\**p*<0.0001. G-J. Bacterial transit across an epithelial barrier was monitored over the course of infection by the genotype I and genotype II clinical isolates characterized above by enumeration of basolateral CFUs. CFUs were calculated 0 m.p.i. (G), 30 m.p.i. (H), 1 h.p.i. (I), and 2 h.p.i. (J). Statistics calculated with a one-way ANOVA. F. TEER was monitored over the course of infection of 16HBE-14o-semi-differentiated monolaters for genotype I and II isolates. B-F. KK03 and KK03ΔΔ*rtxA* are shown as positive and negative controls. All *in vitro* experiments include at least 3, independent biological replicates; averages are plotted with error bars representing the standard error of the mean (±1 SEM).

**Supplementary Tables are attached**.

